# Dynamics of protein binding to sites of nascent unscheduled DNA repair synthesis in non-proliferating cells

**DOI:** 10.1101/2020.03.06.979039

**Authors:** Claudia Scalera, Ilaria Dutto, Francesca Barbazza, Raefa Abou Khouzam, Giulio Ticli, Ornella Cazzalini, Lucia A. Stivala, Ennio Prosperi

## Abstract

The analysis of DNA repair mechanisms is of fundamental importance to understand how cells remove DNA damage and maintain their genome stability. Investigating the dynamic association of proteins at sites of active DNA synthesis has been successfully performed at DNA replication forks, providing important information on the process, and allowing the identification of new players acting at these sites. However, the applicability of these studies to DNA repair events at sites of nascent unscheduled DNA synthesis (UDS) in non-proliferating cells has been never tested. Here, we describe the analysis of dynamics association of protein participating in nucleotide excision repair (NER), and in other DNA repair processes, at sites of nascent UDS in non-proliferating cells, to avoid interference by DNA replication. Labeling with 5-ethynyl-2’-deoxyuridine (EdU) after DNA damage, followed by click reaction to biotinylate these sites, permits the analysis of dynamic association of proteins, such as DNA polymerases δ and κ, as well as PCNA, to active DNA repair synthesis sites. The suitability of this technique to identify new factors present at active UDS sites is illustrated by two examples of proteins previously unknown to participate in the UV-induced DNA repair process.

## INTRODUCTION

Dissecting the mechanisms of DNA repair processes is of fundamental importance to understand how cells remove DNA damage in order to protect genetic information and to maintain genome stability (1,2). However, cancer cells use the same processes to escape chemotherapy applied with DNA damaging drugs. These lines of evidence explain the need of studies to further advance our understanding of mechanistic aspects of DNA repair, with the goal of defining deficiencies in tumor cells which may be exploited for their killing (3).

Among DNA repair systems, nucleotide excision repair (NER) system is an essential DNA repair mechanism able to remove several types of DNA lesions producing helix distortion (4,5). NER alteration is found in rare diseases, such as xeroderma pigmentosum (XP), trichothiodystrophy and Cockayne syndrome (6), and may facilitate tumor formation (7–9). The NER process operates through distinct steps including (i) the recognition of DNA helix distortion; ii) the DNA opening by the pre-incision complex; (iii) the incision and removal of damaged nucleotides, and (iv) the synthesis of the missing DNA fragment, followed by (v) ligation (10). These basic steps rely on a set of proteins whose function has been identified at the molecular level (11). Other studies have investigated the temporal and spatial organization of NER factors (12–14) in order to understand how they are coordinated to promote specific timing and proper localization at DNA repair sites (12–14). These important studies have provided fundamental information on the function of NER factors *in vitro* (11), or on the *in vivo* dynamic behavior of single factors by live cell imaging (15). However, to dissect the mechanistic aspects of the function of this important DNA repair process, and have a complete picture of the temporal and spatial dynamics of the process needs further investigation.

The analysis of dynamic association of proteins with active DNA synthesis sites has been previously shown to provide useful information on DNA replication (16–18), and in particular at functional or damaged DNA replication forks, by means of DNA labeling with 5-ethynyl-2’-deoxyuridine (EdU) (19,20). However, the possibility of applying EdU labeling to nascent DNA repair sites, such as those occurring during unscheduled DNA synthesis (UDS) in non-proliferating cells, has been never tested.

In order to investigate the dynamics of protein operating at active UDS sites, it is very important to exclude significant contribution by the DNA replication machinery, given that many proteins participating in the latter process are also employed in DNA repair. Therefore, the use of normal cells, or non-tumorigenic cell lines that can be rendered quiescent or arrested in the G0/G1 phase of the cell cycle, is imperative for the above purpose. Another important point to be considered is that DNA repair synthesis normally occurs in very short tracts of about 20-30 nucleotides in NER (4,5), making thus challenging not only the number of cells to use for the analysis of protein bound to these sites, but also the choice of optimal conditions (e.g. the extent of DNA damage/lesion, EdU concentration and incorporation time) to enable such type of study.

Here, we have attempted to investigate the dynamic association of protein to nascent UDS sites in growth-arrested cells obtained by serum deprivation or by initial differentiation, such as that occurring in keratinocytes treated with CaCl_2_ (21,22). We show the feasibility of this analysis in normal fibroblasts, and in the HaCaT cell line, and provide evidence that, in addition to known factors involved in DNA repair synthesis, this technique allows the discovery of proteins previously unknown to be present at sites of active UDS after UV-induced DNA damage.

## MATERIAL AND METHODS

### Cell cultures

Immortalized human keratinocytes (HaCaT) were grown in DMEM with high glucose (Euroclone) supplemented with 10% Fetal Bovine Serum (FBS) (Euroclone), 2% Streptomycin/Penicillin (Euroclone) and 2% L-Glutamine (Euroclone).

LF-1 normal human embryonic lung fibroblasts (kindly provided by J. Sedivy, Brown University, Providence; RI, USA) were grown in MEM (Euroclone) supplemented with 10% FBS (GIBCO), 1% Streptomycin/Penicillin (Euroclone) and 1% L-Glutamine (Euroclone). Cells were cultured in sterility conditions and kept at 37 °C in humidified atmosphere at 5% of CO_2_.

Cell quiescence was obtained by serum starvation: LF-1 fibroblasts reached quiescence within 3 days in medium with 0.5% FBS (23), while HaCaT keratinocytes required 5 days in medium with 0.1% FBS.

Calcium chloride (CaCl_2_) (Sigma-Aldrich) was used as supplement in HaCaT cell medium to arrest cell proliferation by inducing cell differentiation (22). CaCl_2_ was added to the culture medium to reach a final 3.8 mM concentration. Cells were grown in CaCl_2_-enriched medium for 5 days.

To evaluate the proliferation state, cells grown on coverslips were incubated for 1 hour in medium containing 20 μM 5-bromo-2’-deoxyuridine (BrdU) or 5-ethynyl-2’-deoxyuridine (EdU), and then cells were fixed in 70% cold ethanol and, if not processed immediately, stored at −20°C.

### Antibodies

The primary and secondary antibodies used are listed in Supplementary Table 1.

### Induction of DNA damage

For UV-C irradiation experiments, cells were exposed to UV light by using a TUV-9W lamp (Philips) emitting at 254 nm. The lamp energy was measured with a radiometer DRC-100X (Spectronics, USA) before every experiment and fixed at 1 J/m^2^/s. Prior to irradiation the culture medium was removed, and cells were washed in PBS. After UV-C exposure, complete medium was added and cells were incubated at 37 °C for the required periods of time.

Local induction of DNA damage was obtained by irradiating cells grown on coverslips through polycarbonate Isopore membranes (Millipore) with 3 μm pores (24).

To induce a DNA lesion repaired by base excision repair (BER), HaCaT cells were treated for 1h with 100 μM 1-methyl-3-nitro-1-nitroso guanidine (MNNG) from a 50 mM stock solution in DMSO. Cells were pre-treated for 15 min with 25 μM O^6^-benzylguanine (BG) to avoid repair by methyl guanine DNA-methyl transferase (25). Both MNNG and BG were from Sigma-Aldrich.

### Setting conditions for optimal EdU incorporation for UDS determination

To set up optimal conditions for determination of proteins bound to nascent UDS sites, EdU incorporation was determined under a range of EdU concentration, incorporation time and extent of DNA damage. To this end, cells grown on coverslips were irradiated as previously described and medium containing different EdU concentrations was added. Cells were recovered at different time points and fixed with cold 70% ethanol and stored at −20°C.

Coverslips were washed with PBS and then click reaction was performed by incubating cells for 30 min with 100 μl of click reaction buffer containing 20 μM biotin azide (ThermoFisher or Jena), 2 mM CuSO_4,_ 10 mM sodium ascorbate in PBS (26,27). A washing step with PBS and then with 1% BSA in PBS plus 0.2% Tween 20 (PBT) was carried out for 15 min. Coverslips were then incubated for 30 min with 50 μl solution containing 1% BSA in PBT and streptavidin-Alexa 488 conjugate (Abcam) diluted 1:150. Samples were washed three times (10 min each) with PBT and then incubated for 1 h with 50 μl solution containing 1% BSA in PBT and biotinylated anti-streptavidin goat antibody (Vector) diluted 1:200. Coverslips were then washed three times with PBT as above, followed by a second incubation for 30 min with 50 μl of streptavidin-Alexa 488 conjugate (28). Afterwards, cells were washed three times with PBT and then incubated for 5 min with a solution of 0.1 μM Hoechst 33258 in PBS, for DNA staining. After two additional washes in PBS, coverslips were mounted in Mowiol with antifading (24).

An Olympus BX51 fluorescence microscope was used for the analysis and significant fields were photographed with Olympus C4040 digital camera. Fluorescence signals of EdU incorporation were then quantified with the Image J software, using the plugins provided. Firstly, the image of nuclear DNA of each considered cell was converted to a 16-bit grayscale image. Then, the Otsu’s function was used to perform automatic image thresholding to identify each nucleus as a single object. After that, the outline of each object was determined via the Analyse Particles plugin and the resulting image was used to create a mask representing the area and the position of each nucleus. Finally, the quantification of the mean fluorescence intensity was measured applying the mask to the corresponding image of EdU staining.

### Isolation of proteins bound to nascent UDS sites

Following initial pilot experiments, we decided to apply a modification consisting in the cell lysis after EdU labeling prior to the cross-linking procedure (29), in order to improve the yield of proteins bound to UDS sites, as described (30).

Cells were seeded at a density of about 70% in 10 cm Petri dishes to reach confluence before proceeding to serum starvation or treatment with CaCl_2_. A total number of 1-3×10^8^ cells were generally used to detect the proteins bound to nascent UDS sites. Complete medium (6 ml/dish) containing 100 μM EdU was added immediately after cell irradiation, or after a period of 10 min, to allow maximal activation of the DNA synthesis step (14). Incubation with EdU lasted not more than 15 min in order to detected proteins associated to nascent UDS sites.

For chase experiments, the EdU was removed and samples were incubated in medium containing 200 μM thymidine (Sigma-Aldrich) for different time points (15 min, 30 min, or 1 h). Then, cells were washed with cold PBS, scraped (two times) on ice using 3 ml of cold lysis buffer (10 mM Tris-HCl pH 8.0, 2.5 mM MgCl_2,_ 0.5% Igepal, 1 mM PMSF, protease inhibitor cocktail), and collected into 15-ml conical tubes. After centrifugation (3 min, 3000 g, 4°C), another step of lysis was performed for HaCaT cells to ensure removal of unbound nuclear and cytoplasmic proteins. The lysis buffer was then removed and cells were washed once with 6 ml lysis buffer without Igepal, followed by washing with 6 ml of isotonic buffer (10 mM Tris-HCl pH 8.0, 150 mM NaCl, 1 mM PMSF, protease inhibitor cocktail). The isotonic buffer was then discarded, and cells were re-suspended with 3 ml of PBS followed by addition of formaldehyde solution to a final 0.5% concentration, to induce the formation of cross-links. The reaction was carried out for 5 minutes at room temperature (RT), and then glycine solution (final 0.125 M) was added to quench cross-linking. After centrifugation (8 min, 3000 g, RT), the supernatant was discarded, and the pellet was stored at −80 °C.

To perform the click reaction, several washes were carried out with PBS + 0.1% BSA. Cells were centrifuged at 4°C for 7 min at 3000 g and the pellets from 5-7 tubes were joined in a single tube. After another wash in PBS, cells were again centrifuged at 4 °C for 7 min at 3000 g and the reaction buffer (20 μM biotin azide, 10 mM sodium ascorbate, 2 mM CuSO_4_ in PBS) was added to the cells (5 ml of click reaction/1×10^8^ cells was used), and tubes were rotated for 1.5 h at RT.

At the end of incubation, samples were centrifuged (3000 g, 7 min, 4 °C) and 5 ml of sonication buffer (50 mM Tris-HCl pH 8.0, 0.2% SDS for LF-1 and 0.5% SDS for HaCaT cells) was used for 1×10^8^ cells, and transferred (about 1.6 ml/tube) into 15 ml-Bioruptor^®^ Plus TPX tubes (Diagenode). Sonication was performed with a Bioruptor^®^ Plus sonication device (Diagenode), setting 30 sec on/30 sec off constant pulse (high power) for 8 cycles. After every cycle of sonication, samples were centrifuged (5000g, 3 min, 4 °C) and the pellet was re-suspended by vortexing. The procedure was repeated four times. From the resulting supernatant lysate, an aliquot was used for protein quantification by the Bradford method; 100 μl were saved as input sample (Inp) and mixed to 3X loading buffer (195 mM Tris-HCl pH 7.4, 300 mM DTT, 30% glycerol, 0.06% blue bromophenol), then boiled for 20 min at 90 °C and stored at −20 °C. To verify the length of DNA fragments after sonication, 30 μl of lysate were taken and 4 μl of 5 M NaCl, 1 μl of 0.5 M EDTA and 1 μl of RNase A (20 mg/ml) were added to the solution. Then, samples were incubated for 30 min at 55°C and after that, 1 μl of proteinase K (20 mg/ml) and 2 μl of 0.5 M EDTA were added for further overnight incubation at 37°C. The DNA fragment size was checked by electrophoresis in TBE buffer on 2% agarose gels (29).

For capture of protein-biotinylated DNA complexes, 100 μl streptavidin-magnetic beads (ThermoFisher or Millipore) were used for 1.0×10^8^ cells. The capture reaction was performed overnight (16-20h) at 4°C on a rotating wheel. Samples were then recovered and centrifuged (3000 g, 7 min, 4°C), the supernatant was carefully discarded, and several washings were performed with PBS + 0.1% BSA on the magnet. The last washing step was carried out with PBS and then 60 μl of 3X loading buffer was added and the samples were boiled at 90°C for 25 minutes and stored at − 20°C. Both input and click capture (CC) samples were then loaded on NuPage 4-12% Bis-Tris gels.

### Western blotting analysis

The protein extraction protocol used to detect the chromatin bound fraction of relevant proteins involved in the NER process has been previously described (24). Briefly, the pellets obtained as described above were resuspended in hypotonic lysis buffer (10 mM Tris-HCl at pH 8.0, 2.5 mM MgCl_2_, 0.5% Igepal, 0.2 mM Na_3_VO_4_, 1 mM dithiothreitol (DTT), 10 mM β-glycerophosphate, 1 mM phenylmethylsulfonyl fluoride (PMSF), protease and phosphatase inhibitors cocktails (Sigma). After washing once in hypotonic buffer, and then in isotonic buffer (10 mM Tris-HCl pH 8.0, 150 mM NaCl, 1 mM PMSF, 1 mM DTT, 10 mM β-glycerophosphate, protease and phosphatase inhibitors cocktails), the pellet was re-suspended in DNase I digestion buffer (10 mM Tris-HCl pH 8.0, 5 mM MgCl_2_, 10 mM NaCl, 1 mM PMSF, protease inhibitor cocktail) and incubated for 20 min at 4°C. Released proteins were mixed in 3X loading buffer, boiled at 75 °C and stored at −20°C. Proteins were separated on NuPAGE 4-12% Bis-Tris gels, and transferred to nitrocellulose membranes for subsequent immunoblot analysis with relevant primary antibodies, and HRP-conjugated secondary antibodies. Nitrocellulose membranes were incubated with substrate Clarity (Bio-Rad) and chemiluminescence detection was performed with a Westar R Chemiluminescence Imager (HiTech Cyanagen) for digital acquisition of images.

### Immunofluorescence analysis

For BrdU detection, cells were incubated with a 2 N HCl solution for 30 min to denature DNA. Afterwards, neutralization was performed with 0.1 M sodium borate (pH 8.0) for 15 min. Cells were then incubated with a blocking solution containing 1% BSA in PBT for 15 min. Coverslips were washed for 5 min with PBT solution and incubated for 1 h with anti-BrdU monoclonal antibody. After washing three times for 10 min each with PBT, coverslips were incubated for 30 min with anti-mouse Alexa 448-conjugated secondary antibody. After that, samples were again washed three times (10 min each) with PBT solution and then incubated 5 min with a solution of Hoechst 33258, as above. Coverslips were mounted with Mowiol containing antifading agent (24). For dual protein detection, coverslips were blocked and incubated with appropriated couple of primary antibodies (mouse and rabbit), following the same procedure described above, and performing the labeling step with anti-mouse (conjugated with Alexa 594), and anti-rabbit (Alexa 488) secondary antibodies.

### Confocal microscopy analysis

Z-stack images have been captured with a Zeiss LSM800 confocal microscope using a Plan-Apochromat 63x, 1.4NA oil-immersion objective (Carl Zeiss). Then, the focal plane showing the maximum fluorescence intensity of green and red channels was selected for the co-localization analysis, which was performed using the freeware ImageJ. In particular, a straight line was drawn in correspondence of the protein fluorescence spot and the peaks of green and red fluorescence intensities along the line were calculated using the Plot Profile function.

### Statistical analysis

Where indicated, the results obtained from immunofluorescence assay were analyzed in order to produce a statistical analysis.

The statistical significance was determined using Student’s t-test using the function of the Prism 6.0 software. A *p-value* lower than 0.05 was considered significant.

## RESULTS

### Setting conditions for EdU incorporation at sites of nascent UDS sites after UV irradiation

In order to detect unscheduled DNA repair in non-proliferating cells, we explored the possibility to use not only primary cell cultures (e.g. fibroblasts) which can be easily rendered quiescent, but also cell lines which may undergo proliferation arrest. The HaCaT cell line of human keratinocytes fulfils this requirement since they are highly proliferating, but can be rendered quiescent by serum starvation, or induced to differentiate by CaCl_2_ treatment (21,22). Thus, residual proliferation was assessed by BrdU or EdU incorporation (Supplementary Figure S1A) either after serum starvation (Supplementary Figure S1B) or CaCl_2_ treatment (Supplementary Figure S1C). We considered acceptable for our purposes the presence of < 5% S-phase cells after 5-day treatment with CaCl_2_, or after serum starvation.

We then sought to set the appropriate conditions of EdU labeling, in order to obtain the maximal sensitivity for detecting proteins associated to short fragments of repair synthesis. To this end, the influence of EdU concentration, as well as the UV irradiation dose on the extent of EdU incorporation were assessed in both LF-1 fibroblasts and HaCaT cells. The results showed that in both cell types, the fluorescence signal obtained with a 10 μM EdU concentration could be increased using a higher concentration, at least up to 100 μM EdU (Figure 1A and Supplementary Figure S1D). A comparable increase in EdU incorporation was found in both cell types by increasing the UV irradiation from 10 to 40 J/m^2^ UV light (Figure 1B). Under maximal conditions (100 μM EdU, 40 J/m^2^), the fluorescence signals increased linearly with time up to at least 180 min (Figure 1C). Therefore, further experiments were performed using these parameters although the EdU incubation time was obviously limited to 15 min, to maintain protein proximity to DNA synthesis sites (16,17).

**Figure 1.**
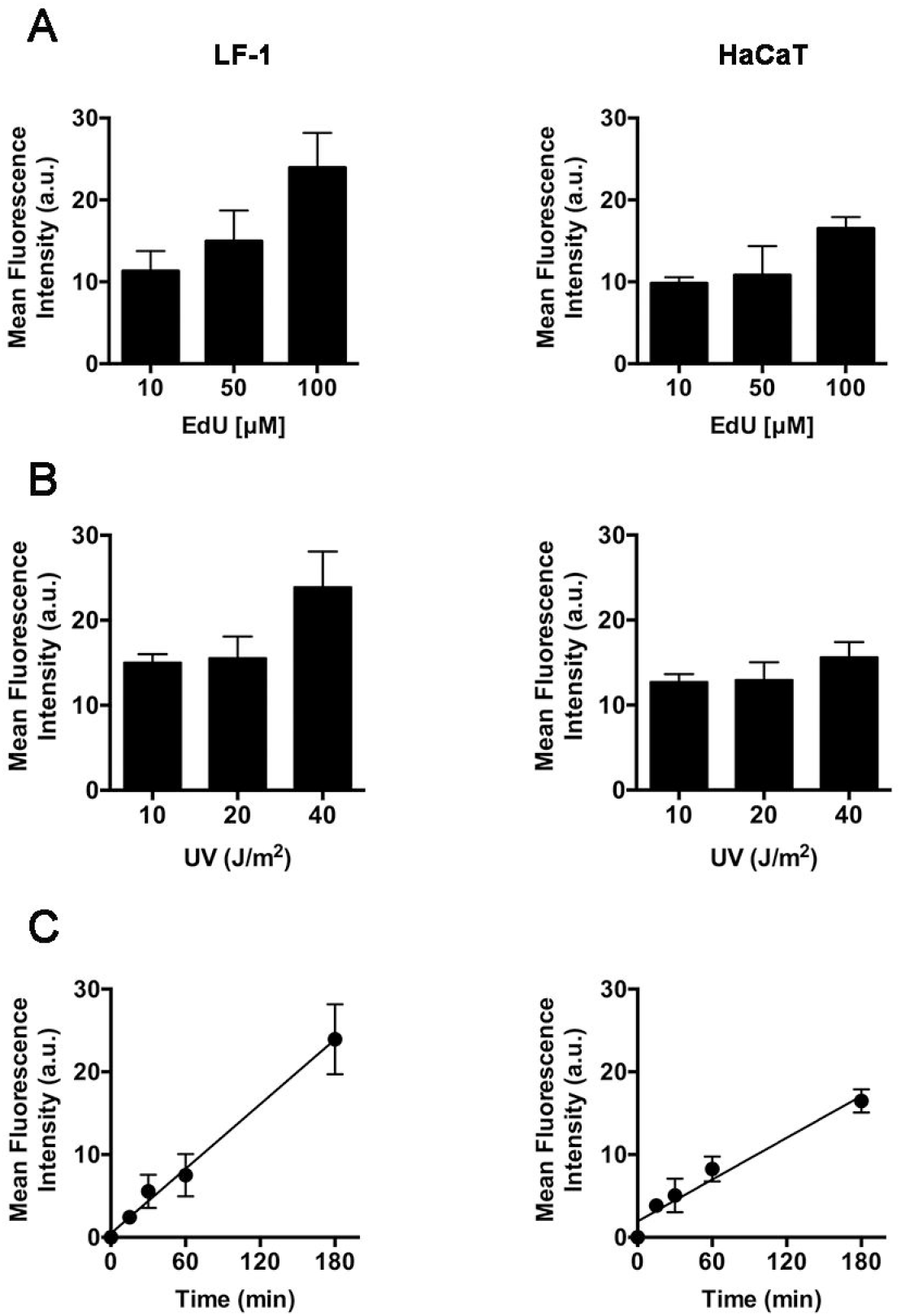
Evaluation of EdU incorporation at sites of nascent UDS sites after UV irradiation. (**A**) LF-1 fibroblasts and HaCaT cells grown on coverslips were growth-arrested by serum deprivation (LF-1), or by CaCl_2_ treatment (HaCaT). After UV irradiation, EdU was added at the indicated concentrations in medium and further incubated for 3 h. Mean fluorescence intensity values of EdU signals were obtained per single cell (*n* > 150 cells per sample, three independent experiments) after image acquisition, and subtraction of background values from cells not exposed to UV. (**B**) LF-1 fibroblasts and HaCaT cells grown as indicated above, were irradiated with the reported doses of UV-C light and then incubated for 3h in medium containing 100 μM EdU. Mean fluorescence intensity values of EdU signals were obtained per single cell (*n* > 150 cells per sample, three independent experiments) after image acquisition, and subtraction of background values from cells not exposed to UV. (**C**) Time course analysis of EdU incorporation in LF-1 fibroblasts and in HaCaT cells grown as described above, exposed to UV-C light (40 J/m^2^), and then incubated in medium containing 100 μM EdU for the indicated periods of time. Mean fluorescence intensity values of EdU signals were obtained per single cell (*n* > 150 cells per sample, three independent experiments) after image acquisition, and subtraction of background values from cells not exposed to UV.

### Analysis of protein capture after EdU labeling of UDS sites after UV irradiation

As the next step, we verified the ability of this procedure to detect typical proteins associated with chromatin after UV irradiation such as PCNA and p125 catalytic subunit (POLD1) of DNA polymerase δ (Supplementary Figure S2). In fact, these proteins were identified after affinity isolation of biotinylated DNA fragments, which were checked to be in the same range applied for iPOND (Figure 2A and Supplementary Figure S2). In particular, densitometry quantification of the amount of PCNA binding to these fragments was significantly higher than in non-irradiated cells (Figure 2B). Similar to HaCaT cells, the procedure was equally efficient in detecting these proteins in LF-1 fibroblasts (Figure 2C). As another specificity control, the isolation of other typical NER factors that are involved in the steps preceding DNA synthesis, (i.e., open complex formation and incision) such as XPA and XPG, respectively, were also tested. The results showed that XPG, but not XPA, could be barely detectable as a factor still associated with on-going unscheduled DNA synthesis (Figure 2D).

**Figure 2.**
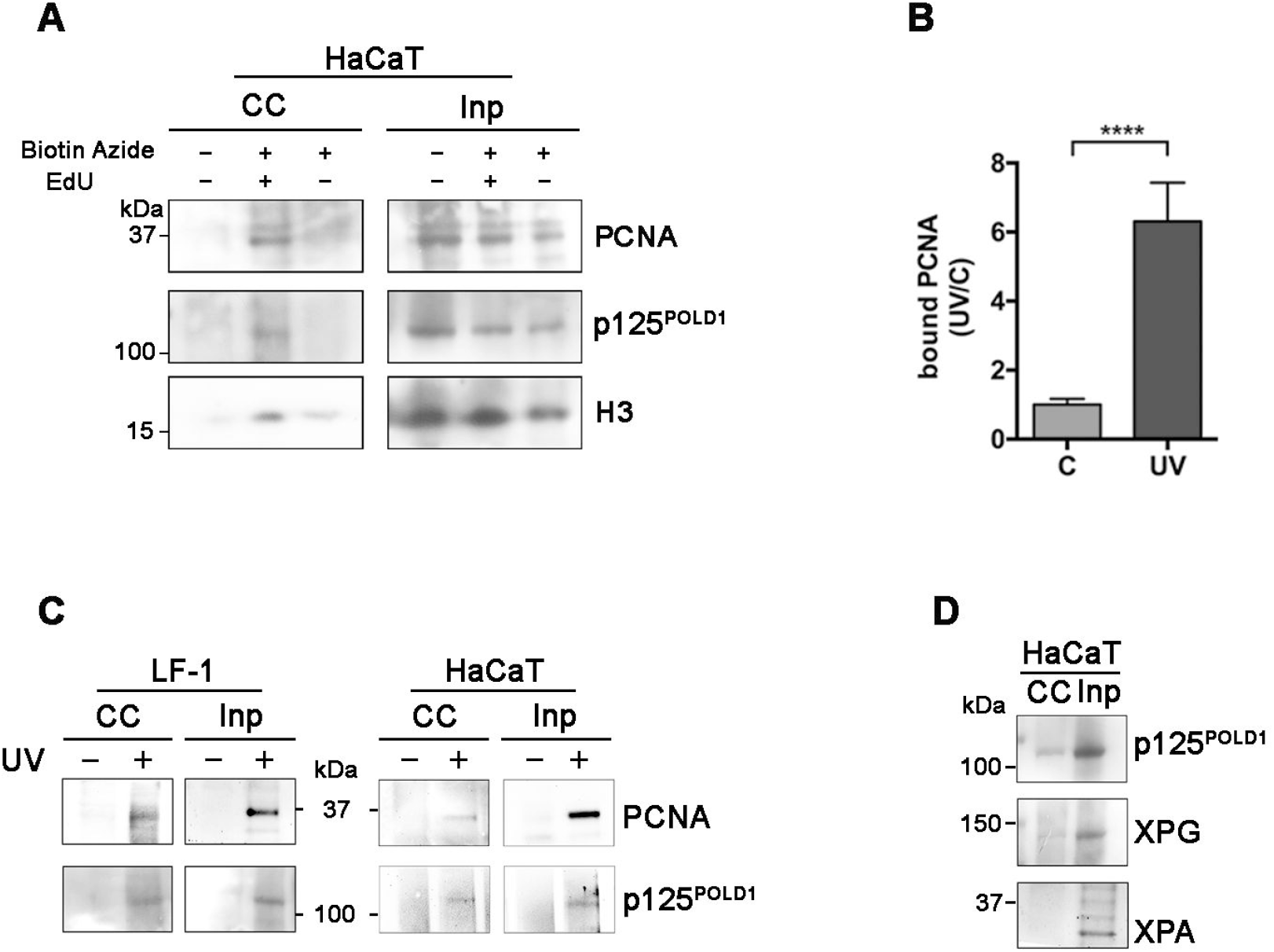
Analysis of protein capture after EdU labeling of UDS sites after UV irradiation. (**A**) HaCaT cells growth-arrested by CaCl_2_ treatment were exposed to UV-C light (40 J/m^2^), and then incubated for 15 min in medium containing (+) or not (−) 100 μM EdU. Click capture (CC) reaction was performed in the presence or in the absence of biotin azide, and captured proteins were analyzed by Western blot for the presence of PCNA, p125 catalytic subunit of DNA polymerase δ (POLD1), and histone H3 (H3). The input (Inp) samples represent loading of 2-4 % of lysate utilized for capture reaction with streptavidin beads. (**B**) Quantitative analysis of the amount of captured PCNA in UV-irradiated *vs* un-irradiated HaCaT cells. Both samples were incubated for 15 min with 100 μM EdU before processing for click capture (CC) reaction. Densitometric quantification of protein signals obtained from Western blot analysis was performed in UV-irradiated cells and the results normalized to that of un-irradiated controls (C). Mean values ± s.d. from 6 independent experiments are shown. (**C**) Comparison of click capture (CC) reaction in UV-irradiated (+) or un-irradiated (−) samples obtained from LF-1 fibroblasts *vs* HaCaT cells. Western blot analysis of captured proteins (CC) and input sample (Inp) was performed for PCNA and the p125 catalytic subunit of DNA polymerase δ (POLD1). (**D**) Click capture (CC) reaction was performed in UV-irradiated HaCaT cells and incubated for 15 min with 100 μM EdU. Western blot analysis of captured proteins (CC) and input sample was performed with antibodies to p125 catalytic subunit of DNA polymerase δ (POLD1), XPG and XPA proteins.

Next, another component of DNA polymerase δ oloenzyme, such as the p66 (POLD3) subunit (Figure 3A), in addition to other typical proteins associated with DNA replication/repair, such as RPA (subunit 2) and CAF1 (p150 and p60 subunits), were identified (Figure 3B). Furthermore, other proteins known to participate in NER in quiescent cells, such as DNA polymerase kappa, XRCC1 and DNA Ligase III (31–33) were detectable, though to different extent, in the EdU-associated fraction (Figure 3C).

**Figure 3.**
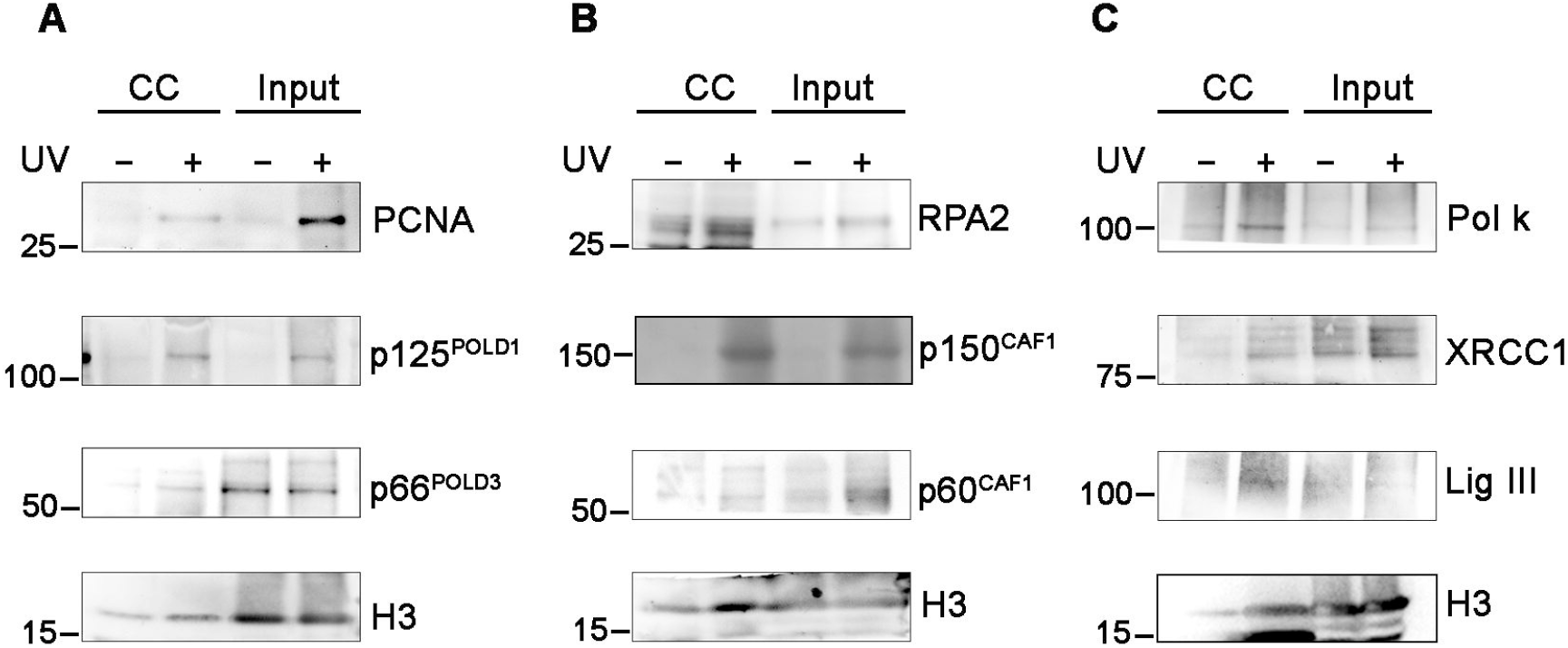
Analysis of NER protein capture after EdU labeling of UDS sites after UV irradiation. **(A)** HaCaT cells growth-arrested by CaCl_2_ treatment were exposed to UV-C light (40 J/m^2^), and then incubated for 15 min in medium containing 100 μM EdU. Western blot analysis of click captured (CC) proteins was performed with antibodies to PCNA and to the POLD1 and POLD3 (p66) subunits of DNA polymerase δ, and histone H3.(**B**) Samples of HaCaT cells grown and treated as in panel A, were analyzed by Western blot for the presence of subunit 2 of RPA protein, the two subunits (p150 and p60) of CAF1 protein, and histone H3.(C) Samples of HaCaT cells treated as above were analyzed by Western blot for the presence of DNA polymerase k (Pol k), XRCC1, DNA ligase III (Lig III) and histone H3.

### Association dynamics of proteins at nascent UDS sites after UV irradiation

Since the association of these proteins to on-going sites of unscheduled DNA synthesis should be limited in time, we verified this behavior (as in the iPOND technique) after a chase period (15 and 30 min) in which excess thymidine replaced EdU to stop its incorporation. The results shown in Figure 4A indicate that, compared with the sample isolated at the end of EdU pulse (0 thy chase), UDS-associated proteins PCNA, p125 (POLD1) and p150 CAF1, were significantly dissociated from the repair sites after 15 and 30 min from EdU pulse. As another feature scarcely investigated in the literature, we wanted to verify whether the association of NER proteins to UDS sites, could be subjected to changes with time during DNA repair. To this end, cells were UV irradiated and then incubated with EdU (15 min pulse) after 15, 60 or 240 min from DNA damage. The isolation of PCNA was detectable at all time points, but other proteins appeared to follow different kinetics (Figure 4B). In fact, p125 POLD1 was detectable only after 15 or 60 min from irradiation, while p66 POLD3 and DNA Ligase III appeared to increase at 4 h after UV exposure, suggesting a more consistent association with late repair sites. This increased association occurred despite the extent of DNA repair was reduced to almost half of the initial rate, as shown by the amount of EdU incorporated during a similar incubation time (30 min) at the indicated periods (0, 60 or 240 min) after UV exposure (Figure 4C and Supplementary Figure S4). This result prompted us to investigate whether the different repair kinetics could be dependent on chromatin accessibility (34,35). To this end, the association of histone H3K9me3 (a typical heterochromatin marker) with UDS sites, was investigated during a similar time course. The results showed that histone H3K9me3 was significantly associated with DNA repair sites at 240 min, as compared with earlier time points, after DNA damage (Figure 4D).

**Figure 4.**
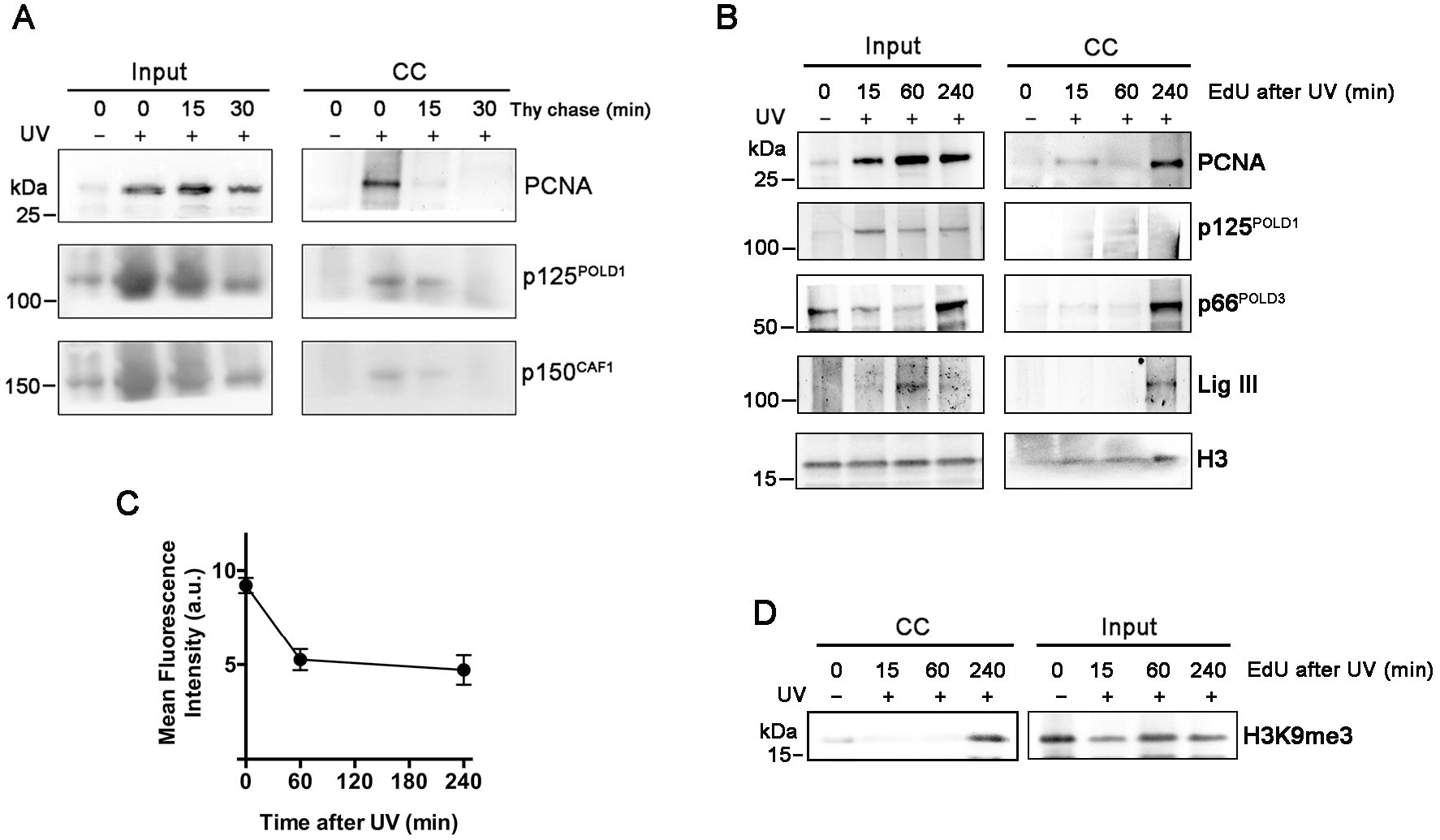
Dynamics of protein binding to nascent UDS sites after UV irradiation. (**A**) HaCaT cells growth-arrested by CaCl_2_ treatment were exposed to UV-C light (40 J/m^2^), then incubated for 15 min in medium containing 100 μM EdU, followed by its removal and chase periods (15 or 30 min) in the presence of 0.2 mM thymidine. Western blot analysis of click capture (CC) *vs* input samples was performed to detect PCNA, the p125 catalytic subunit of DNA polymerase δ (POLD1), and the p150 subunit of CAF1.(**B**) HaCaT cells cultured as in panel A, were UV-irradiated (40 J/m^2^), then incubated for 15 min in medium containing 100 μM EdU at the indicated periods of time after UV exposure. Samples at time point 0 represents cells not exposed to UV. Western blot analysis was performed on CC and input samples to detect PCNA, POLD1 and POLD3 subunits of DNA polymerase δ, DNA Ligase III (Lig III), and histone H3.(**C**) Time course analysis of EdU incorporation in HaCaT cells incubated with 100 μM EdU immediately (time 0), or at the indicated periods of time after UV exposure. Mean fluorescence intensity values of EdU signals were obtained per single cell (*n* > 800 cells per sample, three independent experiments) after image acquisition, and subtraction of background values from cells not exposed to UV. (**D**) HaCaT cells cultured as in panel A, were UV-irradiated (40 J/m^2^), then incubated for 15 min in medium containing 100 μM EdU at the indicated periods of time after UV exposure. Samples at time point 0 represents cells not exposed to UV. Western blot analysis was performed on click capture (CC) and input samples to detect association of histone H3K9me3 to nascent UDS sites.

### Analysis of protein capture after EdU labeling of UDS sites after MNNG treatment

We then asked whether other DNA repair processes could be analyzed with this procedure, and for this purpose HaCaT cells were treated with the alkylating agent MMNG, under conditions known to induce DNA lesions mainly repaired through the BER system (25). Interestingly, major players participating in this process, such as PARP-1, XRCC1 and DNA polymerase β were identified (Figure 5), together with PCNA and p125 (POLD3), which are involved in the long-patch branch of this repair system.

**Figure 5.**
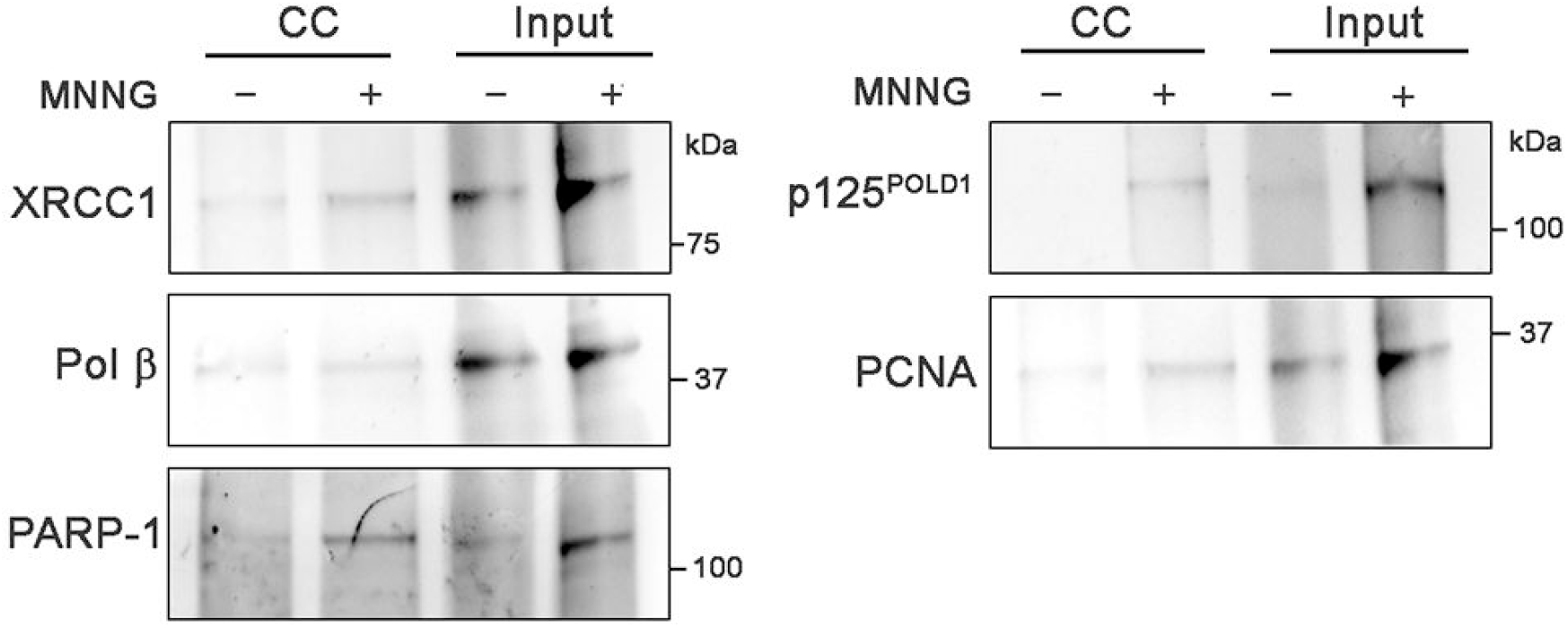
Analysis of protein binding to nascent UDS sites after alkylation DNA damage. (**A**) HaCaT cells growth-arrested by CaCl_2_ treatment were treated with 100 μM MNNG for 30 min and then incubated for 15 min in medium containing 100 μM EdU. Western blot analysis of click capture (CC) *vs* input samples was performed to detect BER proteins such as XRCC1, DNA polymerase β, PARP-1, as well as PCNA and the p125 catalytic subunit of DNA polymerase δ (POLD1).

### Identification of new factors associating to nascent UDS sites

In our hands, the coupling of this technique to subsequent MS analysis, requires a significant amount of starting material, that may be challenging, depending on the type of cells used. In preliminary experiments, we noticed however, that unusual proteins, or factors involved in other processes, such as double strand breaks repair, were captured by this procedure. In particular, two proteins, namely the helicase RUVBL1 and the catalytic subunit of DNA-dependent protein kinase (DNA-PK) attracted our attention. We verified that both factors were recruited to UV-induced DNA lesions, as indicated by immunofluorescence staining and confocal microscopy analysis after local irradiation (Figure 6A, B and D). Interestingly, both proteins were also detected by Western blot after the isolation procedure (Figure 6 C and E), thus confirming the identification by MS of these proteins among those captured by the biotin click capture reaction.

**Figure 6.**
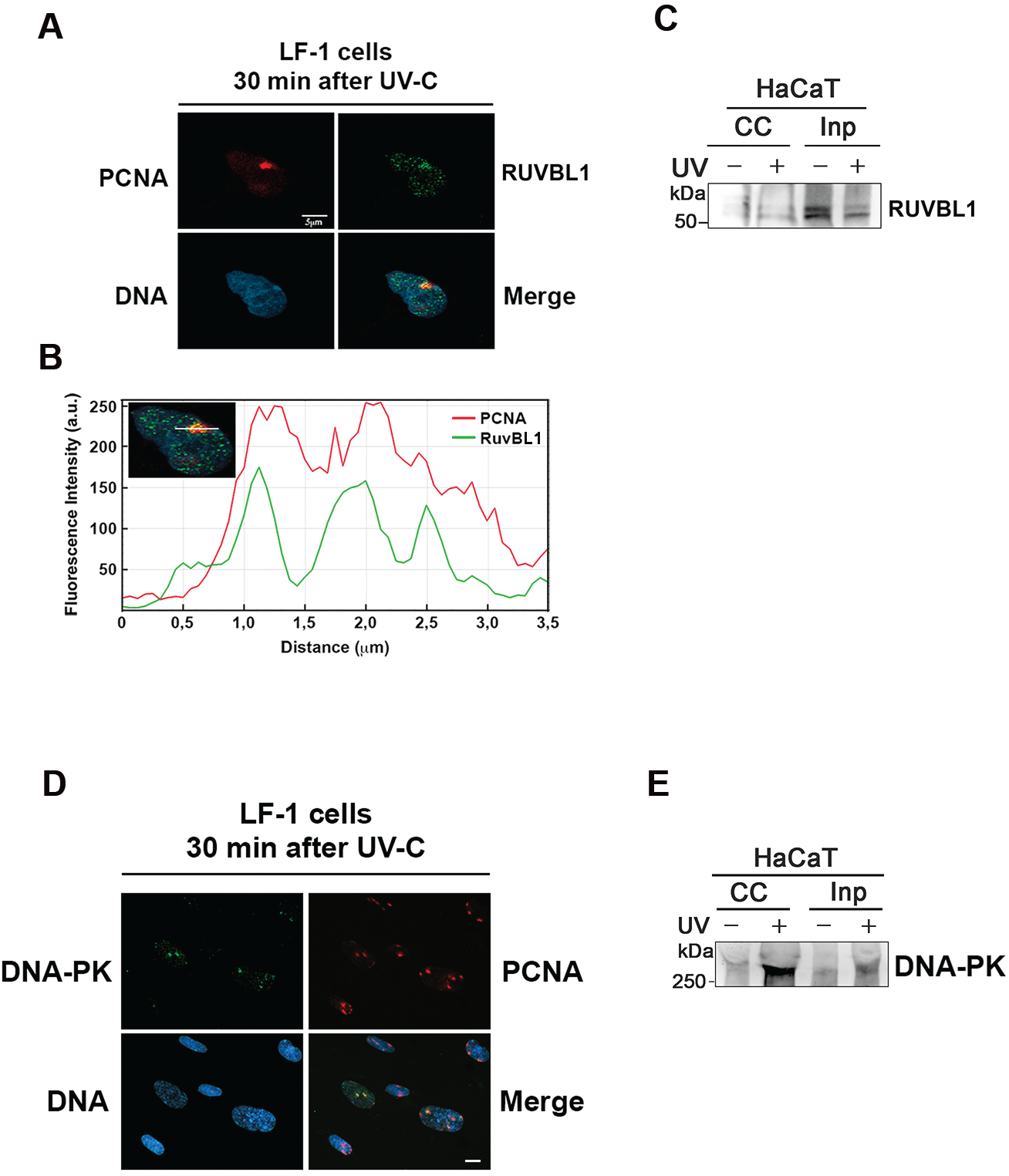
Detection of new proteins binding to sites of nascent UDS after UV irradiation. (**A**) LF-1 fibroblasts were grown on coverslips and growth-arrested by serum starvation. After local UV-irradiation (30 J/m^2^) with Isopore filters with 3 μm pores, samples were lysed *in situ* with hypotonic buffer, fixed, immunostained with antibodies to RUVBL1 (green fluorescence) and PCNA (red fluorescence), and analyzed by confocal fluorescence microscopy. (**B**) Profiles of green and red fluorescence signals along the region defined by the white line shown in the inset. (**C**) HaCaT cells growth-arrested by CaCl_2_ treatment were UV-irradiated as above and incubated for 15 min in medium containing 100 μM EdU. Western blot analysis of captured proteins (CC) and input (Inp) sample was performed with antibody to RUVBL1 protein. (**D**) LF-1 fibroblasts growth-arrested and UV-irradiated locally, as in panel A, were processed for detection of DNA-PK (green fluorescence) and PCNA (red fluorescence); scale bar represent 10 μm. (**E**) HaCaT cells growth-arrested by CaCl_2_ treatment were UV-irradiated and incubated for 15 min in medium containing 100 μM EdU. Western blot analysis of captured proteins (CC) and input (Inp) sample was performed with antibody to DNA-PK protein.

## DISCUSSION

Detecting the dynamics of protein association with nascent DNA at replication forks has been successfully applied with EdU labeling (16,17,19,20). However, this has been not previously investigated at sites of nascent UDS in non-proliferating cells, probably because of the short tract of DNA synthesized during DNA repair (4,5), in comparison with the replication process. Recently, the technique SIRF for *in situ* analysis of protein interaction at replication forks has been developed (28). We have verified whether it could be used to detect proteins associated with nascent UDS in non-proliferating fibroblasts, with no detectable results (not shown). In our study, this limitation could be overcome by using biochemical purification of proteins starting from an amount of cells similar to that used in other techniques, such as iPOND, DmChP, and chromatin capture (16–18). In addition, we have determined conditions that could maximize the amount of EdU incorporation and of proteins associated to UDS sites, to increase the sensitivity of the method. At difference from the above techniques, we have used a permeabilization protocol similar to that used in the accelerated native iPOND, that harvests and lyse cell membranes in a single step, before protein cross-linking, and greatly reduces cell loss that we have experienced following the cross-linking of whole cells (29). Applying these modifications, we have been able to isolate proteins known to be involved in the DNA synthesis steps in non-proliferating cells, such as DNA polymerase δ subunit 1 and 3 (p125 and p66), as well as DNA polymerase k, XRCC1 and DNA ligase III (31–33). However, this is the first time that these proteins have been detected at sites of on-going UDS. Interestingly, among typical NER factors involved in steps preceding DNA synthesis, such as the open complex formation and incision (36), XPG but not XPA could be detected still associated with UDS sites. This finding is in agreement with the evidence that DNA synthesis starts after the first cut performed by ERCC1/XPF complex, while XPG completes the incision after this step (37).

In our study, dynamic binding of these proteins to UDS sites was demonstrated by the thymidine chase experiments, which similarly to the iPOND technique (16) indicated the transient association of the proteins with UDS sites. In addition, we have verified that DNA repair synthesis does not occur with the same rate in the periods following DNA damage. In fact, EdU incorporation was reduced, compared with the initial rate, in agreement with NER kinetics investigated with other parameters, such as chromatin accessibility (34,35). Interestingly, we have found that at time points > 1h after DNA damage, the association of DNA pol δ p125 subunit was reduced, as compared to early times. In contrast, DNA ligase III and the p66 subunit of DNA pol δ, which is known to interact also with other DNA polymerases (38), were more consistently associated with UDS sites at 4 h after UV damage (Figure 3C vs Figure 4B). These findings suggest that the pathway employing other DNA polymerases than DNA pol δ, in conjunction with XRCC1 and DNA ligase III, could operate predominantly at sites of heterochromatin, since the typical marker H3K9me3 was associated to UDS sites at late, but not early time points.

Another important feature of our procedure is that the ability to detect proteins associated with UDS sites during other DNA repair processes, such as BER. In fact, after cell treatment with MNNG, a substance known to induce base alkylation typically removed by BER (25), we could identify the association of DNA pol β, XRCC1 and PARP-1 with nascent UDS sites. Although these factors are involved in the short-patch BER (39), both PCNA and DNA pol δ p125 subunit (long-patch BER) could be also detected, thus indicating that both BER routes were operative.

The possibility to perform proteomic studies by MS identification of new players in the DNA repair process is another challenging aspect that needs further development (19). However, in our initial trials we identified two proteins, DNA-PK and RUVBL1 that were associated with UDS sites, and this was confirmed by their recruitment to local UV irradiation sites. Previous studies indicated that DNA-PK plays a role in NER (40,41). In addition, the activation of ATM signaling during NER was previously shown together with downstream factors MRE11, NBS1, MDC, as well as histone γ-H2AX. Their presence at sites of UV damage was found to be NER-dependent, and attributed to NER reaction intermediates activating the signaling pathway (42,43). Our evidence that also DNA-PK may be localized at active UDS sites is in agreement with these findings.

In contrast, RUVBL1 protein belonging to the family of ATPases with helicase activity, is involved in DNA repair by interacting with chromatin remodeling factors, such as the Tip60/NuA4 and INO80 complexes (44,45). The remodeling function of these complexes in NER has been documented (46,47), although specific involvement of RUVBL1 at DNA repair sites was not provided. Thus, our results indicate for the first time that this protein may participate in the NER process in proximity of UDS sites, although further studies are needed to better clarify the role of this helicase in NER.

In conclusion, our results support the ability of EdU labeling of nascent UDS to isolate factors dynamically associated with DNA repair sites, and open the way to new explorations in this field.

## ACKNOWLEDGEMENTS

We thank prof. J. Sedivy for providing LF-1 fibroblasts.

