## Supplemental data for "Dynamics of protein binding to sites of nascent unscheduled DNA repair synthesis in non-proliferating cells"

Claudia Scalera et al.

### Supplementary Materials

#### Table S1: List of antibodies used in this study

**Primary Antibodies**

| *Protein* | *Company* | *Species/Clone* | *Dilution* |
|---|---|---|---|
| DNA pol δ (p125) | BD Bioscience | Mouse/22 | 1:500 |
| DNA pol δ (p66) | Thermo Scientific | Rabbit | 1:500 |
| DNA pol κ | Abnova | Mouse | 1:400 |
| XRCC1 | Santa Cruz | Rabbit | 1:500 |
| PCNA | Dako | Mouse/PC10 | 1:1000 |
| DNA Ligase III | Genetex | Rabbit | 1:1500 |
| XPG | Sigma | Rabbit | 1:1000 |
| XPB | Thermo Scientific | Rabbit | 1:1000 |
| XPA | Sigma | Rabbit | 1:1000 |
| CAF-1 p150 | Santa Cruz | Mouse/SS11-13 | 1:500 |
| CAF-1 p60 | Bethyl | Rabbit | 1:2000 |
| RPA2 | Bethyl | Rabbit | 1:1000 |
| H3 | Millipore | Rabbit | 1:30000 |
| RuvBL1 | Santa Cruz | Mouse/A11 | 1:2000 |
| DNA PK | Immunological Sciences | Rabbit | 1:1000 |
| Streptavidin | Vector | Goat | 1:200 |
| BrdU | BD Bioscience | Mouse/B44 | 1:50 |

### Supplementary Figures

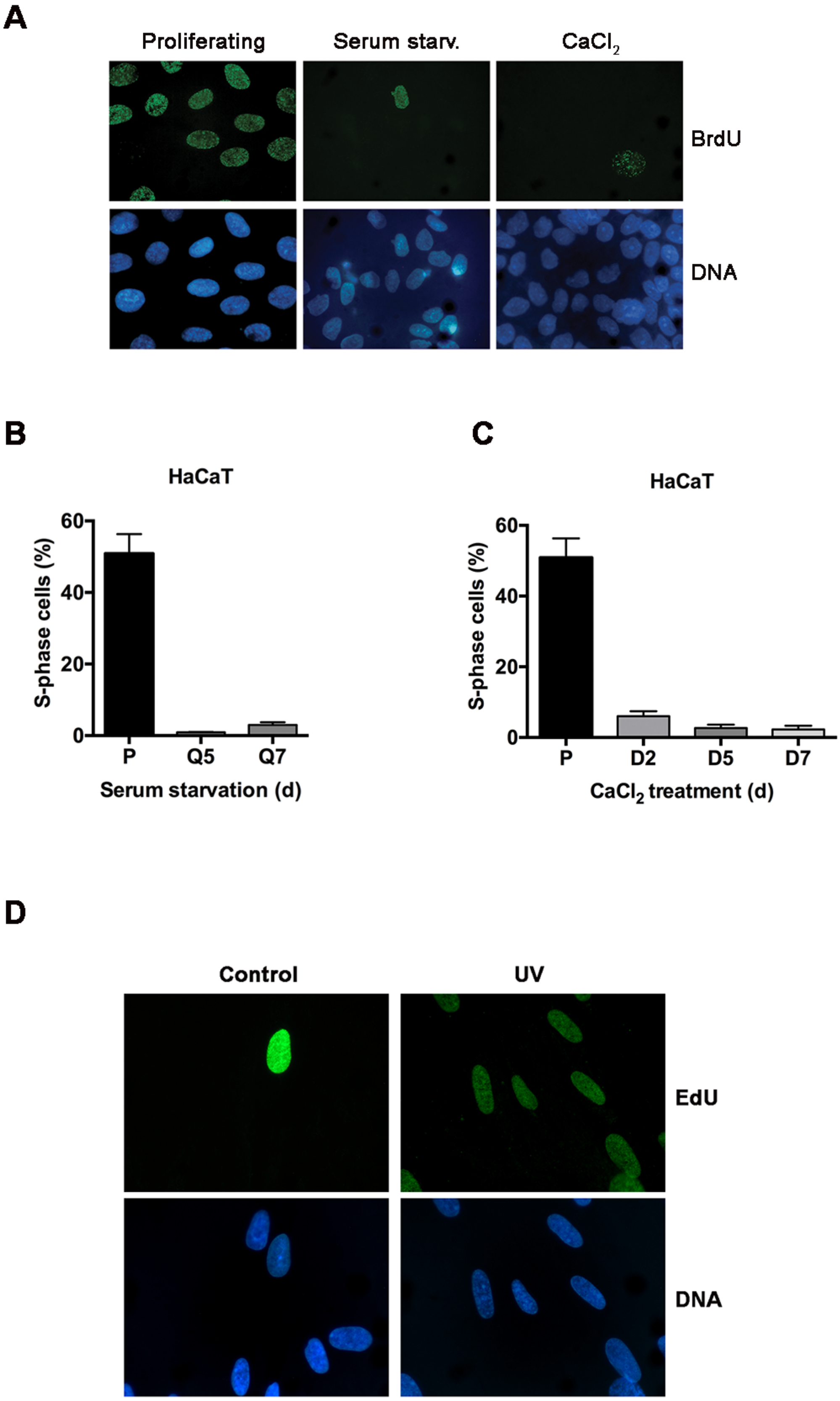

**Figure S1**. Analysis of residual proliferation of HaCaT cells after induction of quiescence by serum starvation, or initial differentiation by $CaCl_2$ treatment. (**A**) Immunostaining of BrdU to mark S-phase cells. (**B**) and (**C**) Quantification of residual proliferation after the indicated days (d) of quiescence (Q) or differentiation (D) induction. P: proliferating cells. Bars report mean values ± s.d. of percentage of BdU-positive cells in three independent experiments. (**D**) Extent of EdU incorporation in untreated (C) and in UV-irradiated (40 $J/m^2$) LF-1 fibroblasts after click reaction with biotin azide and detection with streptavidin-Alexa 488 conjugate. A single S-phase cells is observed in the control cells.

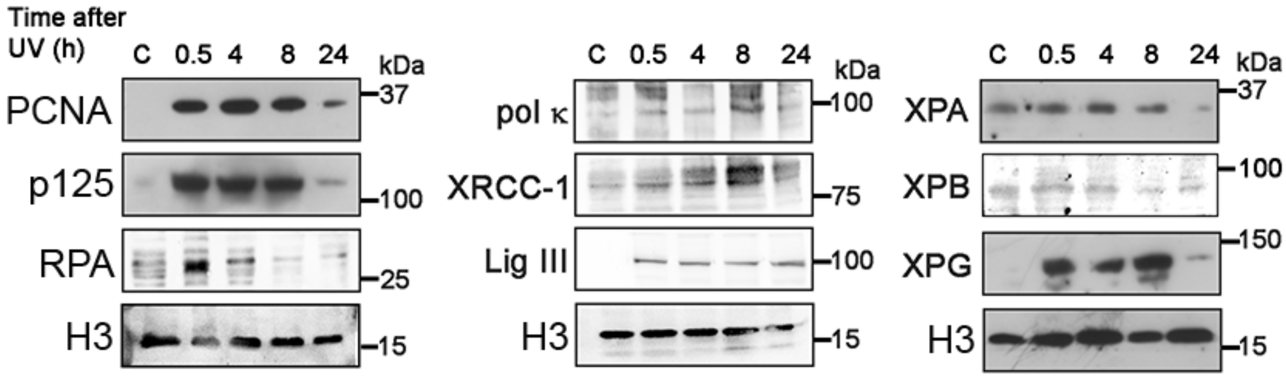


**Figure S2**. Western blot analysis of the recruitment of relevant NER proteins as chromatin-bound form in HaCaT cells, after UV irradiation (10 $J/m^2$) and in non-irradiated control cells (C). Cells were collected at the indicated time points, and chromatin-bound proteins were released as described in Materials and Methods.

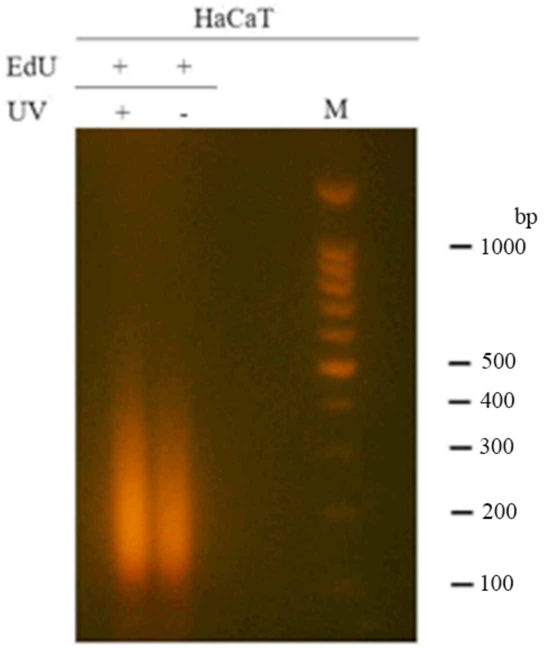


**Figure S3**. Analysis of DNA fragmentation after sonication of samples to be submitted to protein capture with streptavidin magnetic beads.
DNA samples were digested with proteinase K, and RNase A, and analyzed by 2% agarose gel electrophoresis in ethidium-bromide-TBE buffer.

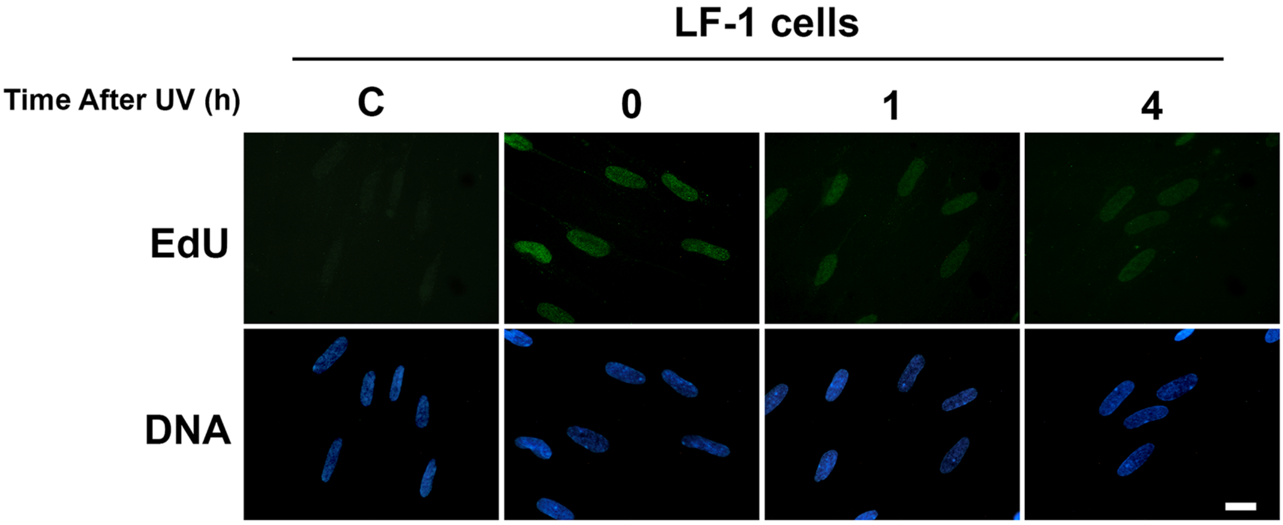


**Figure S4**. EdU incorporation at different periods of time after UV irradiation (40 J/$m^2$). LF-1 fibroblasts growth-arrested by serum starvation were incubated with 100 µM EdU for 30 min, at the indicated periods of time after UV exposure, and then fixed in 70% ethanol. After click reaction with biotin azide, samples were labeled with streptavidin-Alexa 488 conjugate, then with biotinylated anti-streptavidin antibody, followed by a second step of streptavidin-Alexa 488 labeling. Scale bar = 10 µm.
